# An Improved Crystal Violet Assay for Biofilm Quantification in 96-Well Microtitre Plate

**DOI:** 10.1101/100214

**Authors:** Sudhir K. Shukla, T. Subba Rao

**Affiliations:** Biofouling & Biofilm Processes Section, Water & Steam Chemistry Division, BARC Facilities, Kalpakkam, 603 102 India; Homi Bhabha National Institute, Mumbai 400094, India

**Keywords:** Biofilms, Crystal violet assay, Microbial cultivation, 96 well microtitre plates

## Abstract

Microplates are essential tools for biofilm research since it allows high throughput screening of biofilm forming strains or in the assay of anti-biofilm drugs. However, 96 well microtitre plate based assays share the issue of “edge effect”. The primary cause of the “edge effect” phenomenon is evaporation. As edge effect causes a significant increase in plate rejection rate by introducing experimental error, we improvised the classical crystal violet assay to reduce water loss from the peripheral wells. The improvised method showed a significant reduction in edge effect and minimised error in crystal violet assay

## Introduction

Biofilms are surface-attached microbial communities wherein microbial cells are embedded in self-produced extracellular polymeric substances^1^. Biofilms cause severe problems in clinical as well as industrial settings^2^. On the other hand biofilms are used in some of the advantageous applications such as bioremediation of toxic agents and degradation of xenobiotic compounds^3^. In both the cases biofilm production capacity of microbes need to be evaluated in order to use them in bioremediation or to devise strategies against it. Over the last couple of decades, a number of model systems have been tested for *in vitro* study of biofilms^4^. In many assays biofilms are quantified by conventional culture plating method to get colony forming units/count, which is an intensive procedure^1^. Whereas other assays do use 96 well microtiter plates for biofilm quantification as microtitre plate offers comparatively high throughput screening (96 isolates at a time) and it also allows to do screening in multiple sets. Such assays include biofilm biomass assays which are based on staining of matrix including living and dead cells, for example, crystal violet^5^ assay and viability assays that are based on the quantification of only viable cells such as Fluorescein di-acetate assay^6^. On the other hand matrix specific quantification assays involves definite dyes, which bind to the specific matrix components such as living cells, proteins, polysaccharides and eDNA^7^.

A microtiter plate based crystal violet assay is an indirect method of biofilm quantification and was first described by Christensen et al.^8^ Since then several modification are made to increase its accuracy^9,10^. However, micro-titre plate based assays share the issue of “edge effect”. The “edge effect” occurs mainly due to two reasons; first, peripheral wells are more ventilated thus can provide more O_2_ for bacterial growth. Secondly, water evaporates quickly from peripheral wells thereby providing the planktonic cells to stick to the walls, which in turn binds the crystal violet dye and gives a false reading as biofilm biomass. The “edge effect” poses serious concerns when determination of antimicrobial or anti-biofilm efficacy of compounds has to be tested since evaporation increases the concentration of “testing compound” and the experiment end up with wrong crystal violet absorbance values. In this study, some improvisation was made in the crystal violet assay to reduce water loss from the peripheral wells and to reduce the edge effect. The improved method showed a significant reduction in edge effect and minimized the error in crystal violet assay.

## Material and methods

### Microtitre plate assay for biofilm quantification

*Staphylococcus aureus* V329 biofilm was formed on pre-sterilized 96 well flat bottom polystyrene micro-titre plates in triplicates as described elsewhere^11, 12^. Briefly, A 10 μl of cell suspension having 0.5 O.D_600_ was inoculated in 190 μl TSB medium in each well and 200 μl of autoclaved distilled water was added in peripheral wells to reduce the water loss. Then microtitre plate was incubated for 16 h at 37°C. After aspiration of planktonic cells biofilms were fixed with 99% methanol. Plates are washed twice with phosphate buffer saline or sterile saline water and air-dried. Then, 200 μl of crystal violet solution (0.2%) was added to all wells. After 5 min, the excess crystal violet was removed and plates were washed twice and air dried. Finally, the cell bound crystal violet was dissolved in 33% acetic acid. Biofilm growth was monitored in terms of O.D_570_ nm using micro plate reader (Multiskan, Thermo Labsystems).

### Statistical Analysis

All data are expressed as mean standard deviation (SD) of the triplicate experimental data. A two-tailed Student's t-test was used to determine the differences in biofilm formation between the control and each group. The P value of < 0.005 was taken as significant.

### Result and Discussion

Micro-titre plate results showed that when *S. aureus* biofilms are grown in 96 well micro-titre plates for more than 12 h, it is observed that biofilms grown in peripheral wells were thicker (with higher crystal violet absorbance at 570 nm) (Figure 1). CV is a basic dye that binds non-specifically to negatively charged surface molecules such as polysaccharides and eDNA in the extracellular matrix^13^. Because it binds cells as well as matrix components it is generally used to evaluate biofilm biomass *in toto*^14^. However, in practical experience, one needs to take care of some of the experimental artefacts to avoid un-intended errors in our observations/results. As shown in Figure 1, biofilms grown in circumferential wells were grown thicker as compared to biofilms grown in inner wells due to the phenomenon, called as “edge effect”, which primarily occurs mainly due to two reasons; first, peripheral wells are more ventilated thereby provides more O2 for bacterial growth. Secondly, water evaporates quickly from peripheral wells enhancing the planktonic cells to stick to the walls, which bind to CV and give false reading as bioflm biomass. The “edge effect” poses serious concerns when we need to determine antimicrobial efficiency or anti-biofilm efficacy of compounds as due to evaporation concentration of “testing compound” increases and we end up with wrong CV absorbance values. In animal cell cultures experiments, this problem is somewhat managed by using wet incubation chambers that maintains 95% humidity in the incubation chamber. However, normal shaker-cum-incubator which is generally used to grow bacterial cultures does not have this facility. Therefore, to avoid such experimental errors we introduced a simple improvisation in crystal violet method.

**Figure 1:**
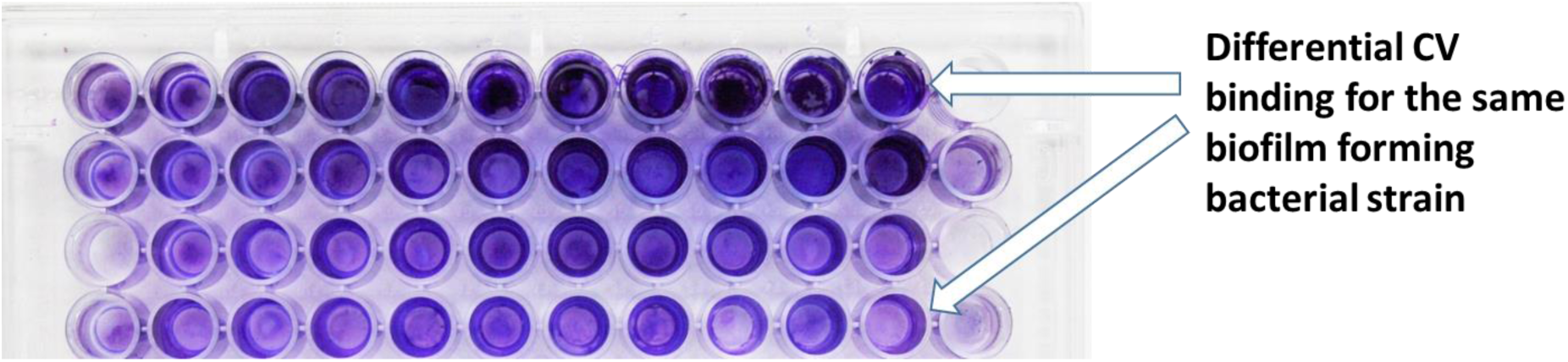
Image showing non-homogenously formed *Staphylococcus aureus* biofilm in 96 well microtitre plate. Apparently peripheral wells have more crystal violet binding to biofilm compared to inner wells.

As said that the primary cause for the “edge effect” phenomenon is evaporation, to reduce excessive water loss and to maintain humidity when we added plain autoclaved water in circumferential wells as indicated in Figure 2. Results showed that adding plain sterile water significantly reduced the edge effect (n=30; p<0.05). As shown in Figure 2, the results showed relative homogeneity in biofilm formation in 96 well microtitre plates with reduced standard deviation in CV absorption values.

**Figure 2.**
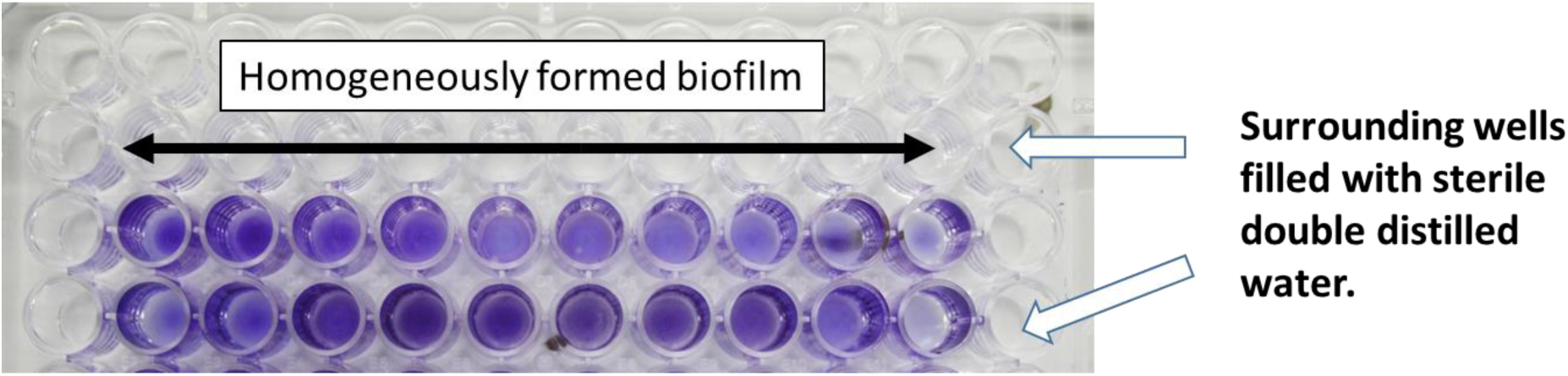
Image showing homogenously formed *Staphylococcus aureus* biofilm when side wells were filled with sterile double distilled water.

In order to keep the time, cost as well as experimental error involved in such microbiological/bioflm assays minimum, it is a prerequisite that the methods used are optimised to produce the best results. Our improvised method shown to give minimum variability in results. This improvisation was highly critical and helpful when biofilm assay was needed to perform with long incubation time involving more than 24 h^15^.

## Conclusion

This improved crystal violet assay method was shown to give minimum variability in our results. This small improvisation in classical crystal violet assay will allow researchers to rapidly screen for biofilm forming bacteria and also to screen anti-biofilm compounds with minimal error and significantly cut down time and money by reducing plate rejection rate.

**Table 1.** Table showing variability in average biofilm growth along the columns in terms of absorbance at 570 nm (mean value ± standard deviation given in parenthesis). First row shows the data for Figure 1 representing the classical method. Second row shows the data for the improved method as shown in Figure 2. *NU = Not used for calculation.

| Method | **Column numbers**<br>(Mean absorbance values ± Standard deviation) | | | | | | | | | | | |
| --- | --- | --- | --- | --- | --- | --- | --- | --- | --- | --- | --- | --- |
|  | 1 | 2 | 3 | 4 | 5 | 6 | 7 | 8 | 9 | 10 | 11 | 12 |
| **Original method** | 0.57<br>($\pm 0.31$) | 0.77<br>($\pm 0.25$) | 1.11<br>($\pm 0.84$) | 1.51<br>($\pm 1.44$) | 1.50<br>($\pm 1.27$) | 1.52<br>($\pm 1.09$) | 1.56<br>($\pm 0.99$) | 1.57<br>($\pm 1.10$) | 1.59<br>($\pm 1.39$) | 1.66<br>($\pm 1.19$) | 1.87<br>($\pm 1.22$) | NU |
| **Improved method** | NU* | 0.96<br>($\pm 0.23$) | 1.22<br>($\pm 0.26$) | 1.05<br>($\pm 0.28$) | 1.24<br>($\pm 0.38$) | 1.12<br>($\pm 0.52$) | 1.29<br>($\pm 0.35$) | 1.33<br>($\pm 0.22$) | 1.26<br>($\pm 0.09$) | 0.98<br>($\pm 0.37$) | 0.86<br>($\pm 0.05$) | NU |

## Conflict of interest

The authors declare no conflict of interest

## References

1. Donlan RM & Costerton JW, Biofilms: survival mechanisms of clinically relevant microorganisms. Clin Microbiol Rev, 15 (2002) 167.

2. Flemming H-C, Biofouling in water systems–cases, causes and countermeasures. Appl Microbiol Biotechnol, 59 (2002) 629.

3. Shukla SK, Mangwani N, Rao TS & Das S, 8 - Biofilm-Mediated Bioremediation of Polycyclic Aromatic Hydrocarbons. *In* Microbial Biodegradation and Bioremediation. Ed., Das S, pp. 203–232, Elsevier, Oxford (2014)

4. McLean RJ, Bates CL, Barnes MB, McGowin CL & Aron GM, Methods of studying biofilms. Microbial biofilms. ASM Press, Washington, DC. 379–413 (2004)

5. Norton T, Thompson R, Pope J, Veltkamp C, Banks B, Howard C & Hawkins S, Using confocal laser scanning microscopy, scanning electron microscopy and phase contrast light microscopy to examine marine biofilms. Aquat Microb Ecol, 16:199–204 (1998)

6. Peeters E, Nelis HJ & Coenye T, Comparison of multiple methods for quantification of microbial biofilms grown in microtiter plates. J Microbiol Methods, 72 (2008) 157.

7. Mangwani N, Shukla SK, Rao TS & Das S, Calcium-mediated modulation of *Pseudomonas mendocina* NR802 biofilm influences the phenanthrene degradation. Colloids Surf B, 114 (2014) 301.

8. Christensen GD, Simpson W, Younger J, Baddour L, Barrett F, Melton D & Beachey E, Adherence of coagulase-negative staphylococci to plastic tissue culture plates: a quantitative model for the adherence of staphylococci to medical devices. J Clin Microbiol, 22 (1985) 996.

9. O’Toole GA & Kolter R, Initiation of biofilm formation in *Pseudomonas fluorescens* WCS365 proceeds via multiple, convergent signalling pathways: a genetic analysis. Mol Microbiol, 28 (1998) 449.

10. Stepanović S, Vuković D, Dakić I, Savić B, Švabić-Vlahović M, A modified microtiter-plate test for quantification of staphylococcal biofilm formation. J Microbiol Methods, 40 (2000) 175.

11. Shukla SK, Rao TS, Effect of calcium on *Staphylococcus aureus* biofilm architecture: a confocal laser scanning microscopic study. Colloids Surf B, 103 (2013) 448.

12. Shukla SK, Rao TS, Calcium-Mediated Modulation of Staphylococcal Bacterial Biofilms. Indian J Geomarine Sci, 43 (2014) 2107.

13. Li X, Yan Z & Xu J, Quantitative variation of biofilms among strains in natural populations of *Candida albicans*. Microbiology, 149(2003) 353.

14. Pitts B, Hamilton MA, Zelver N & Stewart PS, A microtiter-plate screening method for biofilm disinfection and removal. J Microbiol Methods, 54(2003) 269.

15. Shukla SK & Rao TS, Dispersal of Bap-mediated *Staphylococcus aureus* biofilm by proteinase K. J Antibiot, 66 (2013) 55.

